# Extreme phase compression preserves buildable basins in macromolecular crystallography

**DOI:** 10.64898/2026.04.24.720598

**Authors:** Andre LB Ambrosio

## Abstract

Macromolecular crystallography is limited by the phase problem: diffraction experiments measure amplitudes but not the phases required to reconstruct electron density. Existing phasing routes usually seek enough continuous phase information for density modification and model building to converge. Here, we ask how much phase information can be discarded while preserving convergence. We analyzed 14,148 diffraction datasets from chiral crystals to characterize centric reflections in reciprocal-space asymmetric units. After conditioning by centric trace and, where required, index parity, the two theoretical symmetry-allowed phase values were populated near equally, close to 50:50, independent of space group, defining a compact symmetry scaffold. We then retained this exact scaffold while compressing reference acentric phases to a one-bit alphabet {0, π}; as expected from their diffuse parent distribution, the assignments were also near-balanced. Although this binary representation, with fixed attenuation 2⁄π, introduces large angular errors (mean of 52°), it frequently supported automated structure solution: in paired Phenix AutoBuild tests, 705 of 894 binary initializers met a conservative joint criterion of final Free R ≤ 30% and relative chain recovery ≥ 70%, within a 20.0–2.5 Å resolution window. To rank candidate seeds without rebuilding, we developed a branch-balanced Basin Score from inexpensive density-modification and map-connectivity observables computed at 20.0–3.5 Å. The empirical score quickly separates productive from unproductive initializers before AutoBuild. Controlled phase inversion shows that basin compatibility decays gradually and can reappear in an anti-phase-related branch, indicating that buildability is not confined to a single neighborhood around the reference phase set but extends to a much broader field. These results recast phase initialization as basin entry and support future symmetry-aware, binary phase-search strategies.

## INTRODUCTION

Macromolecular crystallography often succeeds even when the initial phase information is imperfect. The key operational requirement is not global correctness of the initial phase set, but whether it positions the map within the ‘basin of attraction’ of iterative real-space and reciprocal-space procedures that can amplify partial correctness into an interpretable structure. This raises a fundamental question: what is the minimal amount of consistent phase information required to ensure entry into this convergent regime?

Detectors measure diffraction intensities at Bragg directions, which are proportional to the structure-factor amplitudes ∣*F*_**h**_∣^2^at reciprocal-lattice points **h**=(*H,K,L*), but not the corresponding phases *ϕ*_**h**_, rendering the direct reconstruction of *ρ*(**r**)an ill-posed problem. The challenge is further exacerbated by imperfect factors such as finite resolution, weak and missing data, anisotropy, bulk solvent, twinning, pseudo-symmetry, and noise (1–4). Classical phasing methods address this by introducing auxiliary constraints to stabilize the inference of *ϕ*_**h**_, such as experimental scatterers, experimental and AI-aided search models, or real-space priors (5–12). In this work, we instead investigate how much the initial phase field can be compressed while still preserving the ability to achieve downstream convergence.

Symmetry divides reflections into centric and acentric classes, which serve as a natural starting point. In the full reciprocal space, a reflection is centric if a space group rotation maps **h** to −**h**, restricting the associated centric phase value 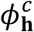 to one of two diametrically opposed values 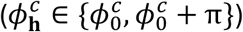, most commonly with 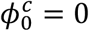or *π*. Translational symmetry elements can introduce origin-dependent offsets, so the allowed phase pair may be shifted (e.g., 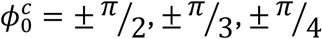), but the two values always remain diametrically opposed. When reducing to the reciprocal-space asymmetric unit (ASU), many of these full-space reversing families collapse onto a small set of observable traces (such as basal, axial, or diagonal), often with simple parity conditions (e.g., even or odd **h** indices or index sums) (13). Acentric reflections lie outside these reversing families, lack any symmetry-fixed phase support; thus, 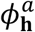 values may populate the circle much more broadly.

Building on this framework, we quantified (i) how restrictive and well-populated the 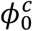 constraints are in practice across macromolecular datasets, and (ii) how aggressively the acentric stream 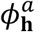 can be discretized while still allowing for successful basin convergence. We found that the most conservative acentric discretization, *K* = 2, where 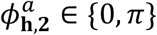, is equivalent to choosing the sign of a real-valued structure factor. When this binary acentric representation is combined with exact two-valued centric supports that are nearly equiprobable, and a global figure of merit (FOM) of 2⁄*π* to account for the overall perturbation, it defines a simple initializer we refer to as attenuated signed amplitudes, or *K*2_*atten*_.

Across a large cohort of macromolecular datasets, we demonstrate that these discrete *K*2_*atten*_ initializers frequently enable successful structure solution using standard density modification and automated model building workflows, such as Phenix AutoBuild (14). Notably, even though the binary phases exhibit a substantial angular discrepancy from the continuous reference phases (with an RMS wrapped error of approximately ∼52°), successful convergence is still the predominant outcome. We further introduce a predictive, low-cost Basin Score calibrated against AutoBuild results, and employ controlled degradation of the phase field to systematically evaluate how reciprocal-space agreement, real-space connectivity, and buildability deteriorate as the binary information is progressively corrupted. Remarkably, even at extreme levels of degradation, convergence can still be achieved for anti-phase sets.

In summary, these results establish three key points. First, exact centric supports combined with one-bit acentric phases are often sufficient for successful basin entry, even when confidence is attenuated. Second, basin proximity can be estimated efficiently enough to guide the search process prior to full model rebuilding. Third, the critical combinatorial factor is not the fraction of binary seeds confined to a single neighborhood around the reference phase set, but rather the fraction that reaches any empirically favorable, rebuildable state, including both solutions similar to the original (control-like) and those related by anti-phase transformation. Thus, *K*2_atten_ serves not merely as an aggressive compression scheme, but as a practical initializer for future basin-guided ab initio phasing, for example, by artificial intelligence, with the Basin Score providing a corresponding soft objective for optimization.

## RESULTS

### Centric burden is universally near-equiprobable across space groups

To isolate a generalizable property of centric phase behavior, we assembled a curated cohort by sampling 15,000 PDB identifiers from an RCSB search restricted to single-chain X-ray structures refined at 1.7 Å or better. For each entry, optimized coordinates and reflection data were retrieved from PDB-REDO when available and converted into standardized per-entry reflection tables. All tables were harmonized to a canonical phase convention on (−*π, π*], annotated with a centric flag, and assigned a new cross-validation set. Unless noted otherwise, all analyses below use a common working resolution window of 20.0–2.5 Å. After validation and filtering, 14,148 datasets spanning all 65 Sohncke space groups represented in the cohort were retained for population analyses.

We define the centric burden of an ASU-reduced reflection table by two coupled descriptors: its extent, measured as the fraction of reflections flagged centric within the working window, and its structure, measured by the number of distinct centric ASU traces and the associated trace-resolved phase supports (Fig. 1). The key empirical result is not simply that centricity collapses into a small number of ASU-observable traces, but that once centric reflections are conditioned on trace and, where required, parity subclass, the two symmetry-allowed phase values are populated at near-equal frequency across datasets, largely independent of space group. To our knowledge, this near-universal trace and-parity-conditioned equiprobability has not been explicitly documented across a cohort of this breadth.

**Figure 1.**
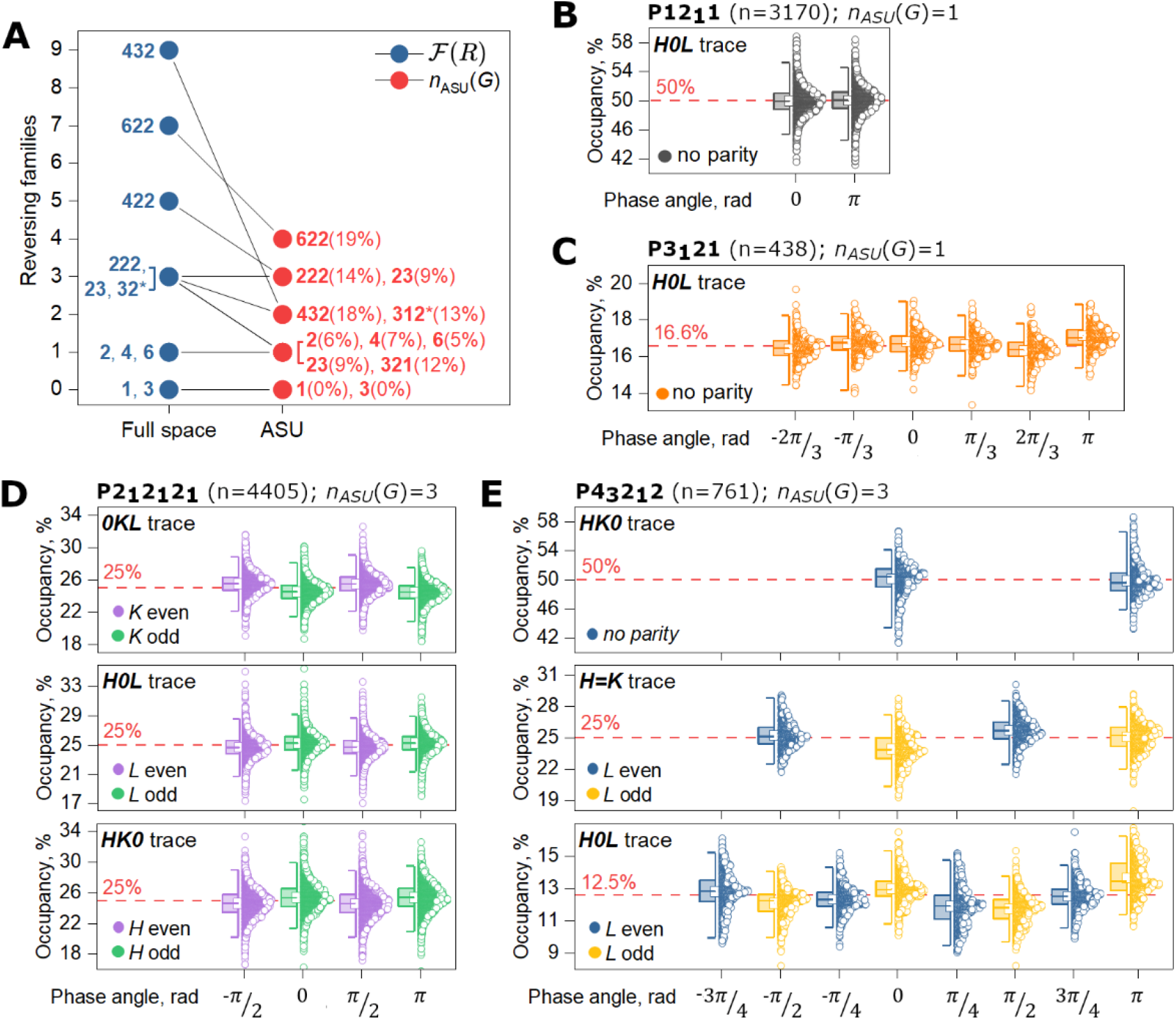
Centric burden collapses into a small set of trace-resolved phase supports in the reciprocal ASU. **(A)** Mapping from the number of reversing families in full reciprocal space to the number of observable ASU traces under the deposited reflection-table conventions. Parenthetical values report the mean centric percentages across datasets for each class. **(B–E)** Representative space groups illustrate how reflection-level diametric supports are expressed after ASU reduction. In P12_1_1, centrics localize to a single trace with the canonical {0,π} support. In P3_1_21, pooled phases span a broader palette because different reflections experience distinct translation-induced offsets, although each reflection still obeys two-point support. In P2_1_2_1_2_1_ and P4_3_2_1_2, parity-conditioned subclasses generate the observable support structure on basal, diagonal, and axial traces. Across all cases, once reflections are conditioned on trace and parity where required, the admissible values are close to equiprobable across datasets. Half-box/jitter plots show per-dataset occupancies; dashed red lines mark the equal-occupancy expectation for the relevant support class. Squares position the means. Whiskers represent the 1^st^ and 99^th^ percentiles.

In full reciprocal space, centricity is carried by a union of plane- and line-like reversing families, *ℱ*(*R*), whose count increases with point-group symmetry: none for point groups 1 and 3; one for 2, 4, and 6; three for 222, 23, and 32; and larger sets for higher-symmetry point groups (Aroyo, 2016). After reduction to the reciprocal space ASU, most of these full-space families collapse onto a much smaller set of observable traces, typically axial, basal, or diagonal. This collapse is practically important because it converts a group-theoretical classification in full reciprocal space into a compact and reproducible description of how centricity appears in deposited reflection tables (Fig. 1A).

The number of centric ASU traces, *n*_ASU_(*G*), provides a compact descriptor of how centricity appears in ASU-reduced data: groups with *n*_ASU_ = 1 typically show low-single-digit centric fractions, whereas groups with multiple traces reach the mid-teens (Fig. 1A). A useful caveat is that *n*_ASU_(*G*) is an ASU-representation property rather than a fundamental invariant. For point group 32, the “321” settings collapse to a single trace under indexing conventions (*n*_ASU_ = 1), whereas “312” settings retain two (*n*_ASU_ = 2), shifting the observed centric fraction without changing the underlying full-space reversing content (Fig. 1A).

The central empirical observation is that, after resolving reflections by trace and, where necessary, by parity or offset subclass, the two admissible phase values are found at nearly equal frequency across datasets. This means that the symmetry-consistent centric prior is not only discrete and exact, but also statistically balanced within each conditioned support. To illustrate this universality, consider several representative space groups. In monoclinic *P*12_1_1, centricity is concentrated on a single ASU trace, the translational offset vanishes on that trace, and the admissible pair reduces to the canonical {0, *π*} support, with near-50:50 occupancy across the cohort (Fig. 1B). In trigonal *P*3_1_21, centricity remains trace-localized, but pooled phases span a six-label palette because different reflections experience different translation-induced offsets; each reflection nevertheless still obeys a two-point constraint, and the pooled labels appear close to uniform (Fig. 1C). In orthorhombic *P*2_1_2_1_2_1_, centricity is distributed across multiple traces and requires parity conditioning; the pooled four-label set decomposes into trace and-parity-specific two-point supports, each again close to equiprobable (Fig. 1D). In tetragonal *P*4_3_2_1_2, screw offsets and parity subclasses produce the richest pooled palette among the examples shown, yet the same principle survives intact: once conditioned on trace and parity, each reflection-level support remains diametric and its two allowed values remain close to balance across datasets (Fig. 1E).

Altogether, group theory fixes the centric scaffold: reversing operations determine which ASU traces and parity subclasses can be centric, and the associated translations fix the corresponding diametric two-point supports. The empirical result is that, once conditioned on trace and parity, these supports are populated at nearly equal frequencies across macromolecular datasets, largely independent of space group, typically within a few percentage points of exact balance. Variation between structures is dominated not by bias within a conditioned support, but by how often the same small set of supports is sampled within the working resolution window. The centric burden is therefore more than a descriptive symmetry constraint: it defines a compact, reusable, and generalizable prior in which the admissible centric phase pair is fixed by symmetry, and the within-support occupancy is effectively equiprobable.

### Binary acentric phases often preserve a buildable initialization channel

Acentric reflections dominate macromolecular diffraction. When the space group contains no reversing families, they essentially comprise the entire phase stream; even when reversing operators are present (*n*_ASU_(*G*)≥1), acentrics usually account for the large majority of reflections (∼80–95 %). Across the cohort, their reference phases are effectively diffuse over the circle: the dataset-wise mean resultant length, 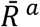, is concentrated near zero, with a median of 0.006 and a 99th percentile near 0.03 (Fig. 2A). Thus, in this regime, acentric initialization is better viewed as controlled discretization than as recovery of a sharply organized prior.

**Figure 2.**
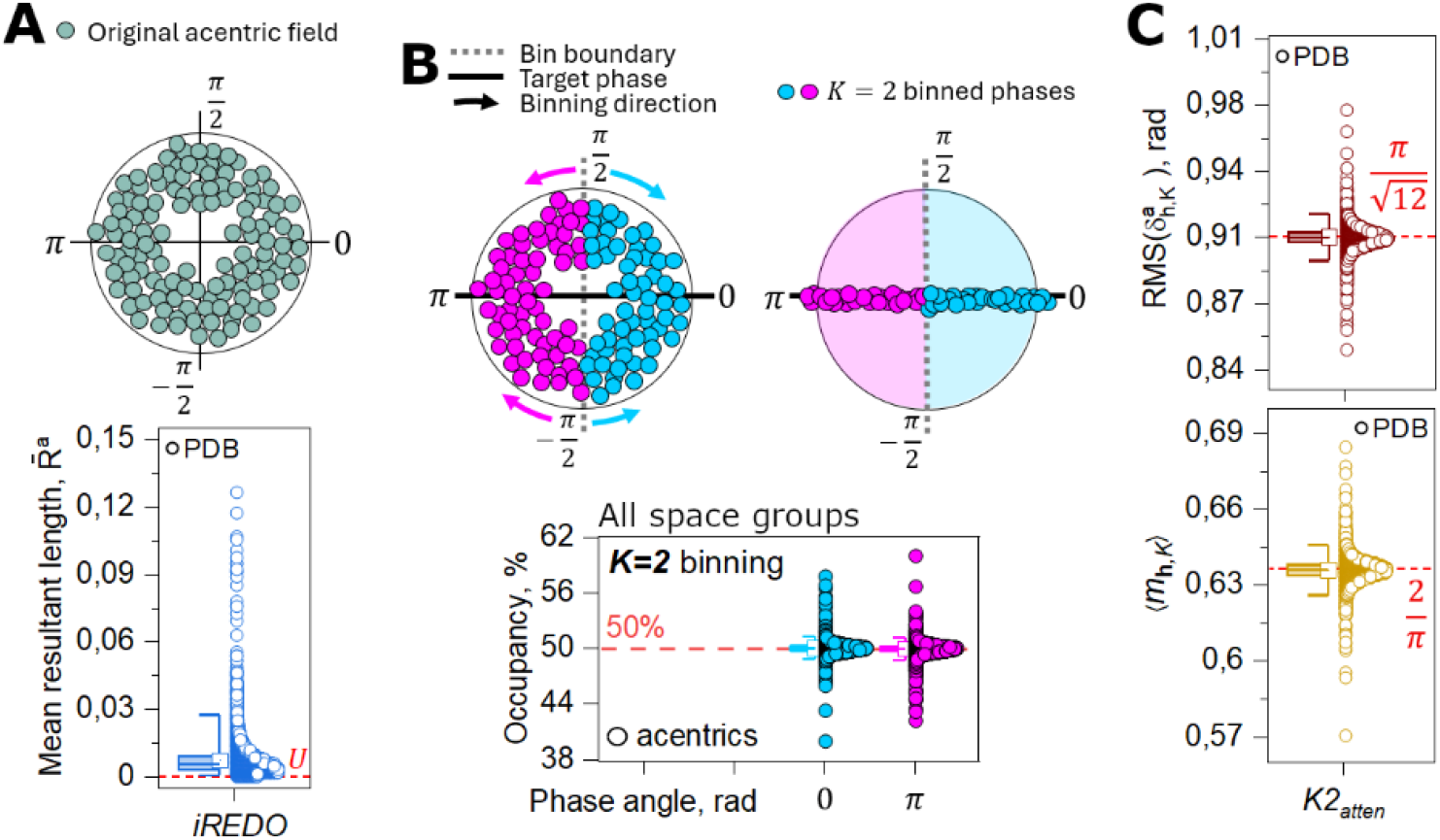
Compression of the acentric phase field. **(A)** Reference acentric phases from the *iREDO* stream populate the unit circle broadly, with low dataset-wise mean resultant length, indicating that the acentric phase field is effectively diffuse near uniformity (U) over the working resolution range rather than concentrated around preferred directions. **(B)** Top: schematic of *K* = 2-acentric binning: each acentric phase is assigned to the nearest of two antipodal targets, reducing the continuous acentric phase stream to a one-bit alphabet {0,*π*}. Bottom: across all space groups, the resulting binary assignments occur at near-equal occupancy, as expected for a broadly distributed parent phase field. **(C)** Empirical consequences of this compression in the cohort. Top: the dataset-wise wrapped RMS angular perturbation induced by *K* = 2 binning closely matches the uniform-phase prediction 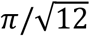. Bottom: the mean cosine attenuation factor, used here as a constant acentric figure-of-merit perturbation prior in the attenuated representation *K*2_atten_, agrees closely with the corresponding expectation 2/*π*. Open circles denote individual datasets; dashed red lines indicate the theoretical values under the uniform-phase approximation. Half-box plots summarize the cohort distributions. Squares position the means. Whiskers represent the 1^st^ and 99^th^ percentiles.

To compress the acentric stream while leaving the centric phases exact, we quantized only the acentric set 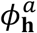 (, CENTRIC=0) onto *K* equally spaced directions on the circle, assigning each phase to its nearest bin center 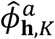. This defines a controlled perturbation where the angular spacing is 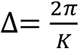 (Supporting Fig. 1); thus, for each *K* binned acentric reflection, the angular perturbation is 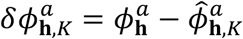. To propagate the resulting loss of information into the standard weighted-map formalism, we attenuated the respective figure of merit (FOM) by the perturbation cosine, 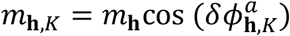.

At the limiting binary compression, *K* = 2, the acentric phase stream is projected onto the antipodal set 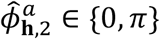. The broad distribution of reference phases yields nearly equiprobable binary assignments, about a 50:50 occupancy across the datasets in the cohort (Fig. 2B). Under this extreme discretization, the empirical acentric 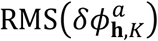 is ∼091 rad (or 52°) per dataset, in close agreement with the rounding-limit prediction of 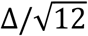 for a uniform error model. The empirical mean FOM attenuation is 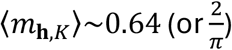, matching the expected mean projection loss under the same uniform approximation of 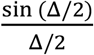 (Fig. 2C). In the conservative 20.0–2.5 Å range isolated from our ≤ 1.7 Å cohort, most reflections exhibit a high signal-to-noise ratio and reference FOM values close to unity; therefore, we applied a uniform attenuated figure of merit and denote the resulting binary acentric representation *K*2_atten_(PHIC_ALL_K2:-FOM_K2_atten), corresponding to a sign-only phase choice for the associated Fourier coefficients, 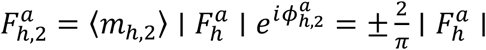.

The central benchmarking result is that this one-bit acentric channel, combined with the exact centric scaffold, frequently remains sufficient for successful downstream phasing (Fig. 3). As an upper-performance reference, initialization from the continuous *iREDO* phase stream gave final AutoBuild Free *R* values centered deep in the buildable regime, with mean 22.8±3.3 %, and median 22.1 %, and 1^st^–99^th^ percentile range 17.5–34.4 % across 925 valid runs. Despite the severe compression of *K*2_atten_, the final Free *R* distribution remained centered in the conventionally interpretable regime, with mean 279+7.6 %, median 25.5 %, and 1^st^–99^th^ percentile range 18.9–52.2 % across 926 valid runs (Fig. 3, left). Only a minority of targets fail to achieve productive density-modification or building trajectories, resulting in poorly refinable maps under standard protocols.

**Figure 3.**
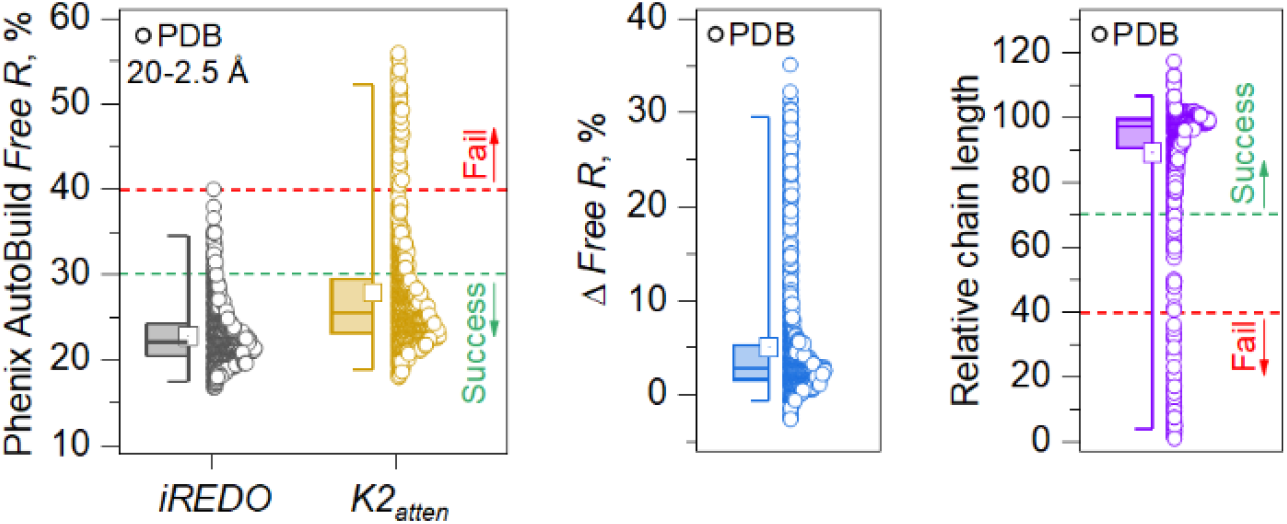
AutoBuild shows that one-bit acentric phases frequently remain inside the convergent basin. Final Phenix AutoBuild outcomes obtained from the continuous *iREDO* initializer and from the compressed *K*2_atten_ initializer over the 20.0–2.5 Å working range. Left: distributions of final Free *R* for *iREDO* and *K*2_atten_. Middle: paired difference in Free *R*, computed as *K*2_atten_ −*iREDO*. Right: chain-level recovery for *K*2_atten_, expressed as residues placed relative to the *iREDO* reference on the paired subset. Despite substantial degradation relative to *iREDO*, the compressed initializer frequently remains within the regime compatible with successful automated rebuilding, consistent with the existence of a broad convergent basin. Dashed lines indicate the conservative strong success and strong failure reference regimes used in the analysis. Half-box/jitter plots summarize per-dataset values. Squares position the means. Whiskers represent the 1^st^ and 99^th^ percentiles. Maximum building and refinement resolution settings were 2.5 Å.

As expected, *iREDO* remained superior, but the loss imposed by binary acentric discretization was usually modest (Fig. 3, center). On the paired subset, *K*2_atten_ increased Free *R* by only 5.1±6.7 percentage points on average (median, 2.8 %; *n* = 925). Chain-level recovery supported the same conclusion (Fig. 3C): relative to the *iREDO* reference, *K*2_atten_ retained a mean chain length of 89.0±21.7 % and a median of 97.1 % on the paired subset with at least 20 residues in the reference build (*n* = 894); 85.8 % of cases retained at least 80 % of the reference chain length, and 20.7 % matched or exceeded it (Fig. 3, right).

Under a joint conservative success criterion used hereon, final Free *R* ≤ 30 % and relative chain length ≥ 70 %, *K*2_atten_ satisfied both criteria simultaneously in 705 of 894 paired cases (78.9 %). These results define a stringent empirical bound on the amount of acentric information required for convergence: global phase accuracy is not the relevant requirement for successful initialization; entry into the basin of attraction of density modification and automated building is. This finding is significant because it establishes a stringent empirical bound on the amount of acentric information required at initialization: while binary compression does not fully solve the phase problem, the benchmark demonstrates that one-bit acentric phases often retain precisely the portion of the phase field that is most critical for successful density modification and automated model building.

### A low-cost Basin Score tracks downstream buildability

AutoBuild provides a direct empirical readout of whether an initializer has entered a productive basin, but it is too expensive to serve as a search objective in a large binary phase space. In practice, individual *K*2_atten_ builds often require more than 40 minutes on a standard personal computer, making direct rebuild-based search infeasible. We therefore sought a cheaper pre-build surrogate that could discriminate productive from unproductive seeds while remaining mechanistically interpretable.

This led us to define a branch-balanced Basin Score, *S*_3.5_, in the narrower 20.0–3.5 Å working window. The score combines two distinct classes of inexpensive observables over a much smaller number of reflections, about a third compared to the 20.0–2.5 Å range. The density-modification branch uses terminal reciprocal-space statistics from *phenix*.*density_modification* (10). The skeleton branch uses coarse real-space descriptors of map connectedness derived from a Clipper-like skeletonization workflow as implemented in Coot (15). Strong success was defined conservatively as final AutoBuild Free *R* ≤ 30 together with relative chain length ≥ 80 %, whereas strong failure required Free *R* > 40 and relative chain length <40 %; intermediate cases were excluded from the primary calibration (Fig. 4). The two branches were modeled separately and then combined with equal weight so that a trajectory could not score highly by appearing favorable in only one domain. Overall, the full set of density-modification and skeletonization metrics, including those not used directly in *S*_3.5_, can be computed together with *S*_3.5_ itself in roughly one-third of a second.

**Fig. 4.**
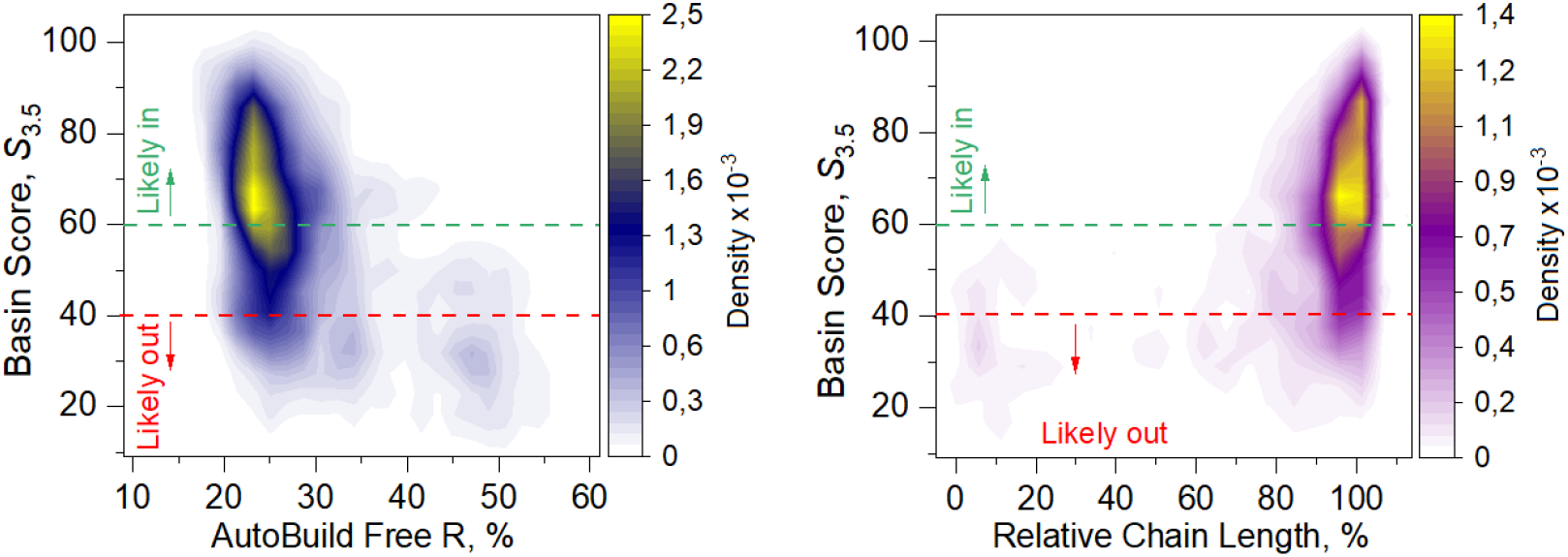
Basin Score distribution with respect to downstream AutoBuild outcome. Two-dimensional density maps showing the distribution of the branch-balanced Basin Score (20.0-3.5 Å working window), *S*_3.5_, as a function of final AutoBuild Free *R* (left) and relative chain length placed with respect to *iREDO* (right) for *K*2_atten_ phase sets. Warmer colors indicate higher point density. Dashed horizontal lines mark the operational score bands used in this work: *S*_3.5_ ≥ 60, likely productive in-basin; 40 ≤ *S*_3.5_ < 60, transition regime; and *S*_3.5_ < 40, likely out-of-basin. The left panel shows that high-scoring trajectories are enriched at lower final Free *R*, whereas the right panel shows that they are likewise enriched at higher relative chain length. Together, these distributions indicate that *S*_3.5_ captures the same productive-versus-unproductive separation inferred from downstream rebuilding, while being computable from inexpensive density-modification and skeleton-connectivity observables evaluated before AutoBuild.

To avoid over-crediting trajectories in which density modification appeared numerically favorable despite a reconstructed map that remained insufficiently protein-like, reciprocal-space and real-space evidence were modeled as separate logistic branches and combined with equal weight. The density-modification branch was defined as *P*_DM_ = σ(−18.072 − 38.171 *R* _FC,FP_ + 34.256 FOM_DM_), where *R* _FC,FP_ is the terminal *F*_*c*_-versus-*F*_*p*_ *R* factor (8, 16) and thus reports reciprocal-space agreement after density modification, and-FOM_DM_ is the terminal mean figure of merit and thus a readout of phase confidence. The skeleton branch was defined as *P*_SK_ = σ(−1.436 + 6.063 LCF_AUC_ − 9.757 EF), where LCF_AUC_ is the area under the largest-component-fraction curve across multiple map thresholds (between 0.8σ and 2.0σ) and thus reports whether the skeleton condenses into a dominant connected protein-like object, whereas EF-is the endpoint fraction at the target map threshold of 1.2σ and thus reports dangling ends and local fragmentation. Here σ(*x*) = (1 + *e*^−*x*^)^−1^. The final score was then defined as *S*_3.5_ = 100(*P*_DM_ +*P*_SK_)/2, such that high values are assigned only to trajectories that simultaneously exhibit reciprocal-space stabilization under density modification and emergence of coherent real-space connectivity.

The resulting score behaves as a practical proxy for basin proximity. High *S*_3.5_ values are concentrated in the region of low final Free *R* and high relative chain recovery, whereas low values are enriched in the converse regime (Fig. 4). Operationally, values *S*_3.5_ ≥ 60 identify likely productive, in-basin states; values *S*_3.5_ < 40 identify likely out-of-basin states; and intermediate values define a transition regime. Thus, the score does more than numerically summarize density modification: it quickly captures the same productive-versus-unproductive separation that becomes explicit only after full rebuilding.

### Basin compatibility erodes gradually and reappears in an anti-phase branch

We next asked how the deployed Basin Score behaves when the *K*2_atten_ phase set is progressively corrupted. To do so, we implemented a controlled-degradation protocol in which an increasing fraction of reflections in the 20.0–3.5 Å scoring window was subjected to phase inversion, *ϕ* ↦ *ϕ* +*π*, and the score was recomputed from the same density-modification and skeleton-connectivity observables. In the representative dataset 2bkf, this window contains 1,737 reflections, of which 506 are centric and 1,231 acentric, and each degradation level in the uniform-random experiment was sampled with 10,000 independent cycles. The undegraded control occupies a strongly favorable region of proxy space (Fig. 5). Under increasing degradation, the mean Basin Score declines steeply, crosses the operational in-basin threshold near 20 % corruption, enters a broad transition regime, and then partially rebounds at the highest degradation levels.

**Fig. 5.**
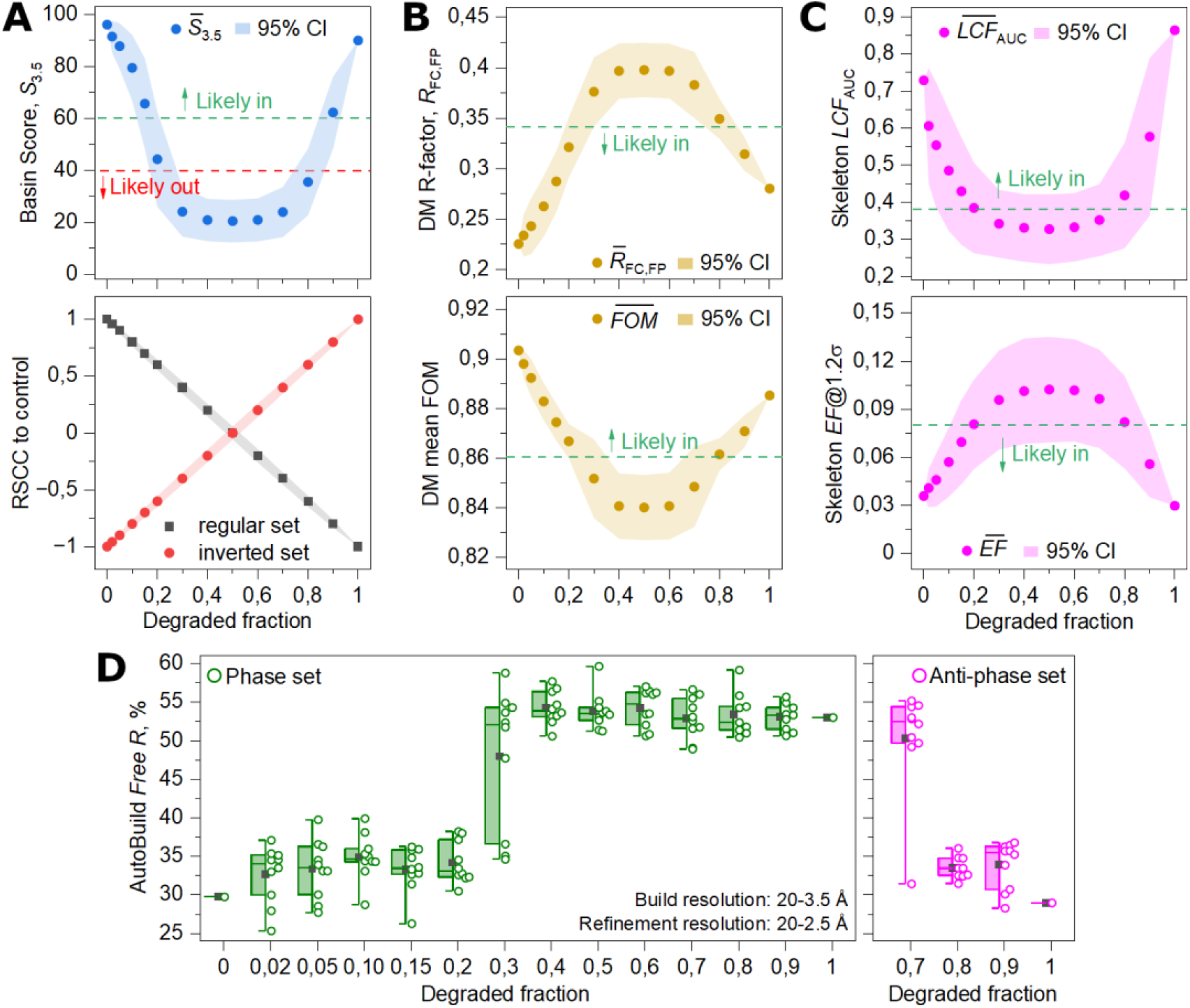
Stability of the deployed Basin Score under controlled degradation of the *K*2_atten_ phase set for PDB *2bkf*. (**A)** Top: mean Basin Score *S*_3.5_ as a function of degraded fraction for uniform-random phase inversion of PDB *2bkf*, with shaded 95 % confidence interval. 10,000 perturbation rounds were performed for each degradation fraction. The undegraded 2bkf control begins at a strongly favorable point, with *S*_3.5_ = 95.97, *R* _FC,FP_ = 0.225, DM mean FOM = 0.903, LCF_AUC_ = 0.729, and endpoint fraction at target sigma *EF*@1.2σ = 0.036. Bottom: signed-control real-space correlations (RSCC) with the regular and inverted control maps. The two curves cross near 50 % degradation, indicating progressive displacement from the original solution toward an anti-control state. **(B)** Top: terminal density modification *R* _FC,FP_ ; bottom: terminal mean FOM, each with shaded 95 % confidence interval and dashed likely-success boundary. (**C), Top**, skeleton LCF_AUC_ ; bottom: endpoint fraction at the target threshold of 1.2σ, again with shaded 95 % confidence interval and likely-success boundary. The degradation series in **(B)** and **(C)** also support conservative empirical working thresholds for productive behavior, approximately *R* _FC,FP_ < 0.34, FOM_DM_ > 0.86, LCF_AUC_ > 0.38, and EF < 0.08. **(D)** Downstream AutoBuild outcome as a function of degraded fraction. For the direct phase set (left), buildability is retained through low degraded fractions and becomes heterogeneous near the basin boundary. For the anti-phase set (right), productive rebuilding reappears at high degraded fractions, approximately mirroring the direct branch at the lowest degraded fractions. Maximum building resolution of 3.5 Å, and refinement resolution settings of 2.5 Å, respectively. This downstream symmetry indicates that controlled corruption can transport the initializer from the control-like branch to a second rebuildable anti-phase branch, expanding the empirically accessible productive basin beyond a single local neighborhood.

The signed-control correlations show that this progression is not merely a gradual accumulation of diffuse noise. Correlation with the undegraded control decreases nearly linearly with degradation, approaches zero near 50 % corruption, and becomes negative thereafter, while correlation with the anti-control rises symmetrically over the same range (Fig. 5A). The degradation series therefore drives the phase set away from the original control branch and toward a distinct anti-phase-related state. The phase trajectory is not simply degrading; it is moving through phase space from one coherent branch to another.

The component metrics clarify how basin exit occurs: the terminal *R* (*F*_*c*_, *F*_*p*_) is among the earliest observables to leave the favorable regime, rising above its likely-success boundary at roughly 20 % degradation (Fig. 5B). The terminal mean FOM remains favorable somewhat longer. In parallel, the largest-component-fraction AUC and the fixed-threshold largest-component fraction deteriorate sharply, whereas the endpoint fraction is more permissive over much of the range (Fig. 5C). Thus, basin loss is driven primarily by breakdown of reciprocal-space agreement and dominant connectedness, rather than by endpoint burden alone. The degradation series also suggests conservative empirical working bounds for productive behavior, approximately *R* (*F*_*c*_ ,*F*_*p*_) < 0.34, DM mean FOM > 0.86, largest-component-fraction AUC > 0.38, and endpoint fraction <0.08. These should be interpreted as practical guideposts rather than replacements for the multivariate score.

Most importantly, the high-degradation rebound is not merely a proxy-level artifact. Downstream AutoBuild confirms renewed buildability in the anti-phase branch (Fig. 5D). The direct phase set remains broadly successful across low degradation fractions and becomes heterogeneous near the basin boundary, whereas anti-phase solutions recover productive rebuilding at high degradation fractions, mirroring the direct branch at low degradation. The degradation series, therefore, reveals not one productive neighborhood, but two empirically accessible rebuilding branches, related by anti-phase transformation and separated by a transition region near half corruption.

This result changes the combinatorial interpretation of binary phase seeding. Although the entire possible solution space for *N* reflections is 2^*N*^ (infinitely small odds for phasing by lucky guessing), the relevant quantity is not the fraction of seeds that reproduce the original control map immediately, but the fraction that enters any empirically favorable, rebuildable state. That favorable region in the K=2 setting includes both control-like and anti-phase-related productive branches. In practical terms, this means that every binary seed should be propagated together with its anti-phase companion.

## DISCUSSION

Our results suggest that, for the tested cohort and conditions, macromolecular phasing can often be approached as a basin-entry problem rather than as a global phase-recovery problem. The initializer does not need to reproduce the full continuous phase field with high angular fidelity. It only needs to place the dataset within a region of phase space where density modification and automated rebuilding can converge. This requirement is often far less demanding than standard intuition would suggest: exact centric supports, combined with one-bit acentric phases under overall attenuated confidence, frequently remain sufficient for productive downstream structure solution, with 705 of 894 paired *K*2_*atten*_ cases meeting the joint success criterion used here.

The conclusion rests on two empirical regularities that sharply reduce the apparent complexity of initialization. First, the centric phase field is not merely symmetry-restricted; it is also statistically minimal. Across 14,148 datasets spanning the chiral space groups represented in the cohort, centric reflections collapse into a small number of ASU-observable traces, and, once conditioned on trace and, where required, parity subclass, the two symmetry-allowed phase values are populated at nearly equal frequency, largely independent of space group. Second, the acentric phase field is effectively diffuse within the working window, so coarse discretization acts as controlled compression rather than destroying a sharply organized prior. In this sense, our approach treats the reduced problem as binary throughout: exact by symmetry for centrics and deliberately compressed for acentrics. The remarkable point is that such a severe representation can still preserve the information required for basin entry in a substantial fraction of cases.

The Basin Score provides a practical handle on that reduced problem. AutoBuild is the definitive downstream assay for determining whether a seed has entered a productive basin, but it is too expensive to use as a search objective. The branch-balanced score developed here shows that basin proximity can instead be estimated from inexpensive reciprocal-space and coarse real-space observables before rebuilding. High values are enriched in the regime of low final Free R and high relative chain recovery, whereas low values are enriched in the converse regime. The score is therefore useful not simply as a descriptive summary, but as a computational objective: it turns the qualitative notion of “close enough to converge” into something that can be ranked, filtered, and optimized in a large discrete search space.

The controlled-degradation analysis reinforces this interpretation in two important ways. First, basin compatibility diminishes gradually rather than collapsing abruptly, which supports the existence of a genuine basin geometry rather than a fragile threshold effect. Second, the degradation series in our tests reveals that productive behavior is not confined to a single neighborhood around the undegraded control. As information corruption increases, the phase set moves away from the control branch, passes through a transition regime of mostly wrong solutions, and then re-enters a productive region associated with an anti-phase-related branch. Downstream AutoBuild confirms that this rebound is not merely a proxy-level artifact, and the practical consequence is immediate: in the binary setting, every seed should be propagated together with its anti-phase companion. The relevant combinatorial quantity is not the fraction of seeds that immediately reproduce the original final map, but the fraction that enters any empirically favorable, rebuildable state.

These results suggest a plausible route for future basin-guided ab initio phasing. The present framework already identifies three ingredients of a viable search strategy: a symmetry-aware reduced initializer, a cheap scalar proxy for basin proximity, and branch-aware downstream validation. Search can therefore proceed not in the original continuous phase space, but in a reduced binary space using moves such as sign flips, block flips, constrained reshuffles, and paired anti-phase transforms, with the Basin Score used as a soft objective to prioritize candidates before full rebuilding. In that setting, the aim is not brute-force enumeration of all binary states, but guided exploration of a reduced space in which a nontrivial fraction of states may already lie inside a productive basin, as observed in our tests. Importantly, being very fast to compute, it makes large stochastic search in a reduced binary phase space, whether by random seeding, iterative sign flips, constrained reshuffles, or related strategies, computationally viable.

Our current implementation is intentionally conservative to serve as proof of principle. The score was calibrated on a restricted cohort of fewer than 1,000 datasets, within a narrow resolution window, using a limited feature set and a simple NumPy/pandas pipeline. Future work should expand this calibration, ideally toward the full PDB-REDO repository. The feature space should grow to include richer reciprocal-space and topology-aware real-space surrogates, including invariant relationships such as centric triplets and quadruplets, threshold persistence, and more explicit measures of map connectedness. The score itself should also evolve, particularly in how it represents anti-control states and basin depth rather than simple basin membership. More efficient implementations, including compiled or GPU-accelerated evaluation, will likely be required for large-scale search to become practical. With the PDB now containing nearly 200,000 crystallographic entries, the scope for large-scale calibration and improvement is considerable.

A particularly attractive future formulation is to unfold all reflection sets to P1 while retaining higher-space-group symmetry as explicit compatibility constraints among binary phase variables. In such a representation, each reflection carries a binary phase state (0 or *π*), while symmetry is enforced through coupling relations rather than through a separate centric/acentric representational split. This would recast initialization as constrained binary optimization over a uniform variable set. The merit of that reformulation is not that it removes symmetry, but that it absorbs symmetry into the search architecture, potentially simplifying proposal generation, scoring, and optimization while preserving crystallographic structure.

At a broader level, the Basin Score may better align the problem with future learning-assisted approaches. The most realistic role for machine learning here is not end-to-end prediction of a continuous phase field, but guided proposal generation in a reduced, symmetry-aware search space: selecting promising seed patterns, prioritizing sign-flip trajectories, or learning basin-proximity surrogates that augment the present score. The significance of the current work is therefore not that it delivers a finished ab initio phasing method, but that it suggests a plausible architecture for one. Exact centric supports, one-bit acentric phases, cheap basin scoring, and systematic branch-aware rebuilding together provide a concrete and empirically grounded starting point for future basin-guided macromolecular phasing.

## METHODS

### Computational organization of the workflow

All analyses were performed with a staged pipeline in which script names encode both the major analysis block and the order of execution within that block. Stage 0 comprised cohort assembly and standardized reflection-table generation, stage 1 quantified empirical centric burden and trace-resolved centric phase structure, stage 2 performed acentric compression and AutoBuild benchmarking, stage 3 calibrated the deployed Basin Score from density-modification and skeleton-connectivity observables, and stage 4 evaluated score stability under controlled degradation together with downstream anti-phase rebuilding tests. Pure Python stages were run with standard Python, whereas Phenix- or cctbx-dependent stages were run under *phenix*.*python* when required. Scripts wrote timestamped logs and structured output folders to facilitate reproducibility and auditability of cohort-scale runs. The entire set of codes, reference files and tutorial, are available on GitHub, at https://github.com/andrelbambrosio/Basin-Guided-Phasing.

### Stage 0: cohort assembly and standardized reflection-table generation

A reproducible cohort of PDB entries was assembled from exported Protein Data Bank query results using *00_select_cohort_ids_with_query_summary*.*py*, which extracted valid PDB identifiers, removed duplicates, and generated a seed-stable randomized subset together with query provenance metadata. For each selected entry, *01_download_build_reflection_tables*.*py* retrieved optimized coordinates and reflection data from PDB-REDO, downloaded FASTA files from the RCSB archive, queried solvent fraction from the RCSB Data API, and generated standardized reciprocal-space CSV reflection tables, employing Gemmi (17) and reciprocalspaceship (18). These tables were harmonized to a canonical PHIC_ALL phase convention (−*π,π*], with −*π* ≡ *π*, annotated with centric flags and cross-validation labels, and supplemented with cell, wavelength, solvent-fraction, and identifier metadata. Optional ECALC-like (19) normalization was performed either per dataset or across the cohort within resolution shells. Space-group composition was then audited with *02_survey_spacegroup_coverage*.*py*, which summarized cohort coverage by space group and crystal system and identified centrosymmetric datasets for exclusion.

### Stage 1: empirical centric-burden analysis

Empirical centric burden was quantified across the cohort using *10_measure_empirical_centric_burden*.*py*, which computed centric and acentric reflection counts, empirical centric phase populations, observed and theoretical unique-HKL centric fractions, and overall and centric completeness. Theoretical counts in the selected resolution window were obtained after reduction to the reciprocal-space asymmetric unit using *Gemmi*. To relate these quantities to symmetry complexity, *11_plot_symmetry_complexity_vs_centric_burden*.*py* computed, for each space group, the number of reversing families in full reciprocal space, the number of ASU-restricted centric traces, and the corresponding point group. Finally, *12_analyze_trace_resolved_centric_phase_supports*.*py* performed a space-group-specific mechanistic analysis by enumerating reversing operations, grouping the associated centric loci into ASU traces, and testing whether each observed centric phase was compatible with the translation-consistent two-point support implied by the corresponding operation. Optional parity-conditioned partitions were used to derive trace-conditioned empirical priors for representative space groups.

### Stage 2: acentric phase compression and AutoBuild benchmarking

Acentric compression was performed only on reflections flagged as acentric in the standardized reflection tables (CENTRIC=0), whereas centric reflections (CENTRIC=1) were retained unchanged. For each requested bin number K, *20_bin_acentric_phases*.*py* quantized acentric phases to the nearest of K evenly spaced angular targets on the unit circle, producing new phase columns and companion attenuated figure-of-merit columns. The script computed dataset-level and global summary statistics for each compression level, including wrapped RMS phase error, mean absolute phase error, upper-tail phase error, attenuation of the figure of merit, and circular-uniformity metrics for the original acentric phase stream. Centrosymmetric space groups were detected and reported separately. Binned reflection tables were written to a derived output directory while the analysis itself remained resume-safe, so that summary statistics could still be generated even when previously binned files were reused.

To benchmark compressed phase initializers in a standard model-building workflow, *21_prepare_autobuild_mtz_inputs*.*py* converted the reflection tables into PHIB-input MTZ files containing amplitudes, amplitude uncertainties, phase estimates, figures of merit, Free R flags, and Hendrickson-Lattman coefficients reconstructed from phase and figure-of-merit values. For the deployed attenuated binary representation, *K*2_*atten*_, the phase source was PHIC_ALL_K2 and the figure of merit was a constant prior K2_ATTEN_FOM = 2/pi when not already present in the input table. HL coefficients were derived from the corresponding circular concentration parameter with a configurable upper cap. By default, this stage sampled a bounded number of datasets using stratified selection across space groups, thereby preserving broad space-group representation rather than allowing highly represented groups to dominate the benchmark set.

AutoBuild benchmarking was performed with *22_run_phenix_autobuild*.*py* using the MTZ files generated in stage 21 and sequence files resolved from the stage-0 download directory. Runs were organized by method and PDB identifier, with one AutoBuild job launched for each method–dataset pair that passed the resume filter. The scheduler was explicitly designed to remain non-idling, maintaining up to a user-defined number of concurrent AutoBuild processes and preferentially completing all requested methods for datasets already in progress before starting new datasets. Successful completion required more than process termination alone: a run was considered valid only if an AutoBuild_run_* subdirectory contained both *overall_best*.*pdb* and *overall_best_refine_data*.*mtz*, and if the corresponding AutoBuild log contained the expected citation marker in its tail. Datasets in which the iREDO phase set produced final coordinates with Free R value > 40 % were deemed as failed and discarded from subsequent analysis.

Downstream AutoBuild results were aggregated with *23_analyze_autobuild_outcomes*.*py*. This stage extracted final Free R values from the overall_best.pdb files, parsed best-cycle residue counts from the AutoBuild logs, and recorded the total number of reflections from the refined MTZ outputs. These results were then summarized both in long format and as a wide per-dataset table containing method-specific metrics. In addition to method-wise Free R distributions, the script computed paired ΔFreeR values relative to a reference method and relative residues placed with respect to the reference build. The latter was used as a pragmatic chain-recovery proxy, with denominator filtering to avoid unstable ratios when the reference build contained very few residues. A compact target table was written for Basin Score calibration, containing the final Free R and relative chain-length proxy for the deployed *K*2_*atten*_ method relative to *iREDO*.

### Stage 3: Basin Score calibration from density modification and map-connectivity surrogates

To derive inexpensive reciprocal-space surrogates of basin proximity, density modification was run independently across multiple resolution windows using *30_run_multiwindow_density_modification*.*py*, a standalone multiwindow wrapper around the Phenix-compatible density-modification helper. The stage accepted either a single reflection table, a list of tables, or a directory of tables, and generated one density-modification job per dataset per resolution window. Default windows spanned 20.0–5.0, 20.0–4.5, 20.0–4.0, 20.0–3.5, 20.0–3.0, and 20.0–2.5 Å, with isolated output subfolders for each window. Each job was executed in an independent operating-system process with hard wall-clock timeout enforcement, allowing stalled runs to be terminated safely without corrupting the broader batch. Parsed log summaries reported for each dataset and window the number of reflections used, terminal density-modification mean FOM, terminal R-factor between Fc and Fp, the probability-map correlation, and the bias ratio when available.

To derive inexpensive real-space surrogates of basin proximity, *31_reconstruct_maps_and_extract_skeleton_proxies*.*py* reconstructed CCP4 maps directly from the reflection tables for one or more specified phase:FOM pairs. For each pair, the script generated both a raw map from F exp(iφ) and a FOM-weighted map using Gemmi for MTZ construction and Fourier transformation. Skeletonization was then performed on either the raw or FOM-weighted map after thresholding, with threshold values interpreted in sigma, absolute-density, or percentile units. The skeleton graph was built with configurable voxel connectivity and optional recursive tip pruning, after which threshold-dependent connectivity metrics were computed (map σ values of 0.8, 1.0, 1.2, 1.4, 1.7, and 2.0), including the number of connected components, size and fraction of the largest connected component, endpoint fraction, branchpoint fraction, mean degree, cyclomatic number, and related quantities. Threshold-integrated quantities, including the area under the largest-component-fraction curve, and fixed-threshold quantities at a user-specified target threshold (map σ=1.2), were summarized per dataset and phase:FOM pair.

We used ChatGPT 5.4 as an analytical aid to evaluate the density-modification and skeleton-proxy summary tables produced in this workflow (steps 30_ and 31_). Based on these inputs, it assisted in selecting the two most predictive metrics from each branch, estimating the associated logistic-model coefficients, and comparing candidate models across the full set of resolution windows for prediction of Phenix AutoBuild success and failure (steps 22_ and 23_). The equal-weight model adopted in the manuscript, S_3.5, was obtained through this ChatGPT 5.4-assisted model-selection procedure.

### Stage 4: controlled Basin Score degradation and downstream rebuilding

The stability of the deployed Basin Score was evaluated by controlled corruption of the binary initializer using *40_BasinScore_degradation*.*py*. This stage took a binned reflection table as input and treated the PHIC_ALL_K2 phase set, together with its attenuated figure-of-merit prior, as the control state. Within the selected scoring window, a user-defined fraction of reflections was subjected to 180° phase inversion, with independent handling of acentric and centric subsets according to the reflection classification in the input table. Three degradation modes were implemented: uniform-random flipping, low-to-high resolution flipping, and amplitude-descending flipping. For each degradation fraction and each replicate round, the script generated a degraded phase set, computed real-space correlation to both the undegraded control and the sign-inverted control, and then re-evaluated the low-cost reciprocal-space and real-space basin-proximity signals used in the deployed score. The stage was designed for large numbers of replicate trials, with process-level isolation, timeout protection, chunked execution, checkpointing, and resume support.

An important feature of this stage is that it did not reimplement all downstream operations directly inside the main driver. Instead, it orchestrated a set of dedicated helper APIs for density modification, skeleton analysis, and MTZ export. The thin stage-40 wrapper *40_api_density_modification*.*py* provided a stable entry point for density modification and delegated the actual calculation to *phenix_like_density_modification_from_csv_py27*.*py*, which reconstructed CCTBX (20) arrays from the reflection-table CSV, generated or reused Hendrickson-Lattman coefficients, and ran the RESOLVE/Phenix density-modification engine in a manner designed to mimic *phenix*.*density_modification* (10) as closely as possible under *phenix*.*python*. Real-space scoring relied on a two-layer skeleton-analysis interface: the batch engine 40_api_reconstruct_map_and_skeleton_batch.py reconstructed maps, thresholded the selected map representation, built a skeleton graph, and computed the same connectivity metrics used in stage 3, while the bridge wrapper 40_api_skeleton_proxy.py loaded that batch engine dynamically, constructed the required argument namespace, ran a single CSV state, and emitted a compact JSON summary of the proxy metrics. AutoBuild-ready degraded states were written by *40_api_autobuild_mtz_writer*.*py*, which reconstructed Hendrickson-Lattman coefficients from PHIB and FOM and wrote PHIB-input MTZ files with the columns required by AutoBuild.

For each degraded state, reciprocal-space and real-space proxy metrics were recomputed and combined into the same branch-balanced Basin Score logic used in the deployed stage-3 model. The degradation controller therefore recorded, for every task, the terminal density-modification R(Fc,Fp), terminal mean FOM, largest-component-fraction AUC, endpoint fraction at the target threshold, and the resulting Basin Score. It also explicitly tracked whether a degraded state passed or failed the individual component thresholds associated with the productive regime defined in the deployed score. Real-space correlation to the control and anti-control maps was evaluated in parallel as an independent diagnostic of displacement from the undegraded solution and approach to the anti-phase-related branch. Sampled degraded states could also be exported as CCP4 maps and AutoBuild-ready MTZs, with manifests written incrementally as degradation fractions were completed.

To test whether highly degraded states could re-enter a productive branch related by anti-phase transformation, companion MTZ files were generated for exported degradation states above a user-defined minimum degradation fraction using *41_make_antiphase_degradation_mtzs*.*py*. The anti-phase generation script recursively scanned the degradation output tree, located PHIB-input MTZs associated with selected degradation fractions, and constructed anti-phase companions by applying PHIB ← PHIB + 180° mod 360° and sign inversion of the first two Hendrickson-Lattman coefficients (HLA, HLB), while leaving HLC and HLD unchanged. These seed and anti-phase states were then benchmarked by downstream *phenix*.*autobuild* with *42_run_phenix_autobuild_from_degradation_mtz*.*py*, which consumed manifest CSVs produced by the degradation workflow, launched matched seed and anti-phase AutoBuild runs in separate work directories, and applied the same strict run-validity criteria used in the stage-2 AutoBuild benchmark. Finally, *43_analyze_autobuild_degradation_results*.*py* pooled outcomes across degradation fractions, degradation modes, and phase variants, and assembled paired seed-versus-anti summaries by matching runs that shared the same degradation mode, degradation fraction, and round number. These outputs were then used to quantify anti-minus-seed differences in Free R, residues placed, and success fraction across the degradation series.

## ACKNOWLEDGEMENTS

This work was supported by the São Paulo Research Foundation (FAPESP) through grants 2021/05726-6, 2013/07600-3, and 2024/04805-8. The author is grateful to Mateus Piovezan Otto and Felipe Souza Lincoln, and to FAPESP scholarships 2018/23675-7 (to M.P.O.) and 2018/23946-0 (to F.S.L.), for valuable early collaboration, discussions, critical feedback, and exploratory code contributions that helped shape the computational analyses developed in this study. The author thanks Professors Glaucius Oliva and Alessandro Silva Nascimento (IFSC/USP) for insightful discussions, suggestions, and feedback throughout this project. The author also thanks IFSC/USP for technical, computational, and infrastructural support.

